# Cholesterol transport cycle of human ABCA2

**DOI:** 10.64898/2026.08.25.746927

**Authors:** Shi Min Tan, Kilian Schnelle, Natalia Voskoboynikova, Mateusz Nowacki, Bianca M. Esch, Florian Fröhlich, Michael Holtmanspötter, Jacob Piehler, Dmitry Sharev, Kristian Parey, Dovile Januliene, Arne Moeller

**Author notes:** <u>For correspondence:</u> (A. M.).

## Abstract

Cholesterol is a key component of cellular membranes and is critical for brain function, particularly axon myelination. Among the 48 human ATP-binding cassette (ABC) transporters, ABCA2 exhibits the highest expression in the brain and is involved in cholesterol metabolism, primarily in oligodendrocytes. Notably, ABCA2 has been associated with myelin sheath integrity and maintenance, as well as Alzheimer’s disease. Here, we report cryo-EM structures of human ABCA2 that reveal critical endogenous lipid-binding sites unique to ABCA2. Our five distinct conformations include a previously uncharacterized intermediate between the closed and apo states of ABCA subfamily transporters. Most importantly, we elucidated the cholesterol transport mechanism of ABCA2, which involves novel interdependent rotations of the exocytoplasmic domains (ECDs) and regulatory domains (RDs). Our structural findings provide a new perspective on ABCA transporter function and highlight the role of ABCA2 in facilitating efficient cholesterol recycling and transport in the brain.

## Introduction

Cholesterol is a primary lipid component of human cellular membranes and a major constituent of myelin sheaths in the brain^1,2^. Myelin membranes, derived from the plasma membrane of oligodendrocytes, ensheath and insulate neuronal axons to enable rapid impulse conduction^1,3^. Dysregulation of cholesterol homeostasis in the central nervous system (CNS) is associated with major neurological disorders, such as Alzheimer’s disease, Huntington’s disease, and Niemann-Pick C disease^4^. This association underscores the critical role of tightly controlled cholesterol regulation in the brain^2^. Although the brain comprises only 2-5% of total body mass, it contains approximately 25% of the body’s unesterified cholesterol^2,4^. Only a small proportion of cholesterol is degraded, secreted, and synthesized in the brain, and the blood-brain barrier separates it from peripheral tissues, preventing cholesterol uptake from the bloodstream^5^. Furthermore, brain cholesterol exhibits a long half-life, lasting up to five years, compared to only a few days for peripheral cholesterol^2,4^. Accordingly, brain cholesterol homeostasis requires specialized protein machinery within the endolysosomal pathway to ensure efficient cholesterol recycling and trafficking within the brain^6^.

Lipid transport within the CNS is mediated by several human ATP-binding cassette (ABC) transporters. Among these, the ABCA subfamily is essential for maintaining cellular lipid homeostasis^7^. Mutations in these transporters are linked to various genetic diseases that arise from disrupted lipid homeostasis^8,9^. For instance, ABCA1 mutations cause Tangier disease, ABCA3 mutations are linked to fatal neonatal pulmonary surfactant deficiency, and ABCA4 mutations result in Stargardt disease^9^. ABCA2 is the most abundant ABC transporter identified in both human and rodent brains^10,11^. It is predominantly expressed in oligodendrocytes and, to a lesser extent, in neurons^12,13^. Previous studies established ABCA2’s involvement in lipid transport and cholesterol metabolism within myelinating cells of the CNS, contributing significantly to the integrity and maintenance of the myelin sheath^14,15^. Dysregulation of ABCA2-mediated sterol transport is implicated in several human disorders, including Alzheimer’s disease and cancer^16^. As such, an ABCA2 polymorphism is associated with both early- and late-onset Alzheimer’s disease^17,18^. Notably, ABCA2 is also implicated in chemotherapy resistance^19,20^. A fivefold increase in ABCA2 expression has been observed in ovarian carcinoma cells, conferring resistance to estramustine^21^. Elevated ABCA2 expression has also been reported in several human brain cancers, including vestibular Schwannomas^19,20^.

Like other ABC transporters, members of the ABCA subfamily consist of two homologous halves, each comprising a transmembrane domain (TMD) and a nucleotide-binding domain (NBD)^7^. Hallmarks of ABCA transporters are their large dimeric exocytoplasmic domains (ECDs) and two regulatory domains (RDs), which engage in domain swapping to facilitate interactions with the opposing NBD^7^. With these additional components, the ABCA subfamily comprises the largest members among the seven ABC subfamilies^22^. Although ABCA transporters share structural similarities, the function of each transporter varies with cellular localization, tissue-specific expression, and substrate specificity^23^. While most characterized ABCA transporters mediate lipid transport across the plasma membrane, ABCA2 is mainly associated with intracellular cholesterol and lipid trafficking within endosomal and lysosomal compartments^24^. Initially identified as an endolysosomal transporter involved in cholesterol efflux and membrane lipid dynamics, ABCA2 now appears to have broader functions beyond lipid transport and regulation of membrane lipid homeostasis^23,24^.

Despite substantial progress in understanding ABCA2 and ABCA family transporters, many fundamental questions regarding their molecular mechanisms and function remain unresolved. Specifically, the processes by which lipids are extracted from the membrane and transported to and through the ECDs are not fully understood. Furthermore, the intricate interplay among RDs, NBDs, TMDs, and ECDs, as well as their conformational changes driving the transport cycle, requires further elucidation.

Here, we performed structural characterization of ABCA2 in different states using cryo-electron microscopy (cryo-EM). By combining delipidation and relipidation approaches, we resolved three nucleotide-free conformations and two ADP-bound post-hydrolysis conformations at resolutions between 2.6 Å and 3.4 Å, which jointly elucidate the mechanism of cholesterol transport. Furthermore, we confirmed the cellular localization of ABCA2 using lysosomal pull-down mass spectrometry and defined the orientation of ABCA2 within the cellular organelles using dual-color confocal fluorescence microscopy and super-resolution DNA-PAINT. Our findings delineate ABCA2’s previously reported roles in cholesterol homeostasis within oligodendrocytes and in myelin sheath formation, and highlight several unique features of this enigmatic ABC transporter.

## Methodology

### Expression and purification of ABCA2

Human ABCA2 gene obtained from Genscript (Uniprot: Q9BZC7 isoform 3) with a C-terminal Flag tag in pcDNA3.1 vector was expressed in HEK293S GnTI^-^ cells (ATCC) by transient transfection. Cells were harvested after 48 hours post-transfection and lysed using a dounce homogenizer in 25 mM HEPES pH 7.4, 200 mM NaCl, 20% glycerol, 2 mM MgCl_2_ and cOmplete EDTA free protease inhibitor cocktail (Roche) before solubilizing with either a mixture of 1% Lauryl Maltose Neopentyl Glycol (LMNG) and 0.1% cholesteryl hemisuccinate (CHS) (Anatrace) or 1% LMNG for 2 hours at 4°C. The solubilized lysate was centrifuged at 100,000 x *g* for 1 h, and the supernatant was incubated with anti-FLAG M2 affinity resin (Sigma Aldrich). The resin was washed with 20 column volumes (CV) wash buffer (25 mM HEPES pH 7.4, 200 mM NaCl, 5% glycerol and 0.02% LMNG-CHS/LMNG) before eluting with 5 CV of elution buffer (25 mM HEPES pH 7.4, 200 mM NaCl, 5% glycerol, 0.002% LMNG-CHS (or 0.002% LMNG) and 100 μg/ml 3X FLAG peptide). The elution fraction was concentrated and applied to size-exclusion chromatography, using Superose 6 Increase 3.2/300 column (Cytiva), equilibrated in 25 mM HEPES pH 7.4, 200 mM NaCl, 0.001% LMNG and/or LMNG/CHS. The peak fractions were collected for cryo-EM sample preparation and ATPase activity assay.

### ATPase activity assay

Pyruvate kinase/lactate dehydrogenase-coupled ATPase assays were performed as previously described^25^. Briefly, the enzyme reactions were assayed in a 96-well plate with a total reaction volume of 200 μL. Purified ABCA2 (10 ug) was added to the ATPase assay reaction buffer (50 mM Tris, pH 8, 12 mM MgSO_4_, 0.3 mM NADH disodium salt, 0.05 mg/ml pyruvate kinase, 0.05 mg/ml L-lactic dehydrogenase, 0.1 mM EGTA, 3 mM phosphoenolpyruvate and 10 mM ATP). Absorbance at 340 nm was recorded every two minutes at 37°C for 2 hours using a TECAN Infinite M Nano+ plate reader to monitor NADH oxidation.

### Cryo-EM sample preparation and data acquisition

For cryo-EM specimen preparation, purified ABCA2 was concentrated to 2.5 mg/ml. For vanadate-trapped post-state, 5 mM Na_3_VO_4_, 10 mM ATP and 20 mM MgCl_2_ were added to the purified ABCA2 sample and incubated at 37°C for 30 mins. 2.5 μL of sample was applied onto the glow-discharged UltrAuFoil R1.2/1.3 gold grids (Quantifoil) and vitrified using Leica EM GP 2 with >80% humidity at 4°C. The grids were blotted for 6 sec before plunge freezing in liquid ethane. Cryo-EM data were acquired at 200 kV with a Thermo Fisher Glacios electron microscope and a Falcon 4i camera with a Selectris energy filter at a nominal magnification of 165k. The slit width was 10 eV and aberration-free image shift was used during data collection. The calibrated pixel size was 0.68 Å/pixel and total exposure was set to 50 e-/Å2 at a defocus range of -1.8 to -0.6 μm. All three datasets were collected in Electron Event Representation (EER) mode using the EPU data collection software v2.9 (Thermo Fisher).

### Image processing

Cryo-EM datasets of the pre-states and the vanadate-trapped closed- and return state of ABCA2 were processed in cryoSPARC (v.5). The overall workflow is summarized in Supplementary Fig. 2. Movie frames were corrected for beam-induced motion using patch-based motion correction, and contrast transfer function (CTF) parameters were estimated using patch-based CTF estimation in cryoSPARC live (v.5). Micrographs were filtered by CTF fit resolution using a 5 Å cutoff, yielding 20,415 micrographs for the pre-state dataset, 19,692 for dataset containing pre-state up and down conformations and 20,857 micrographs for the dataset containing closed- and return-states. Particles were picked using blob-based, template-based and proprietary in-house picking methods and extracted with a box size of 512 pixels. They were Fourier cropped to 128 pixels and 2D-classified separately per picking method. Selected particles were merged and duplicates were removed after a further round of 2D-classification. This resulted in stacks of 4,147,960 particles for the pre-state, 1,736,418 for the pre-state up and down and 875,015 particles for the closed- and return-states. Selected particles were used to generate a single ab-initio 3D reconstruction, while unselected particles were used to generate five decoy models using ab-initio 3D reconstruction with five classes. These six models served as input for one round of heterogenous refinement to remove junk particles and the remaining particles were used to generate new ab-initio 3D reconstructions with five classes. These initial models were then subjected to three rounds of heterogeneous refinement to remove low-quality particles and separate distinct conformational states where present. Particles were re-extracted with a box size of 672 pixels and Fourier cropped to 512 pixels. Multiple rounds of non-uniform (NU) refinement combined with orientation rebalancing, subset selection based on per-particle scale and global CTF-refinement and finally reference-based motion correction gave final stacks of 237,977 for the pre-state, 174,265 for the pre-state up, 248,291 for the pre-state down, 101,256 for the closed-state and 106,755 for the return-state. Final non-uniform and local refinements with C1 symmetry yielded reconstructions at 3.37 Å for the pre-state, 2.64 Å for the pre-state up and 2.63 Å for the pre-state down, while the vanadate-trapped closed-state reached 2.90 Å and the return-state 2.87 Å.

### Model building and refinement

Initial atomic models were generated using the AlphaFold3 prediction^26^, which was placed in the individual maps and fitted as rigid bodies through ChimeraX^27^. The structure was manually inspected in Coot^28^ and iteratively refined using phenix.real_space_refine in combination with rigid-body refinement within PHENIX^29^. Validation reports were automatically generated by MolProbity^30^ indicating stereochemistry with 0.00 and 0.16% outliers (all-atom clashscore: 5.84 and 7.93). Refinement and validation statistics are summarized in Table 1. Figures were generated in ChimeraX^27^.

### Dual color confocal fluorescence microscopy

HeLa WT cells were seeded in 35 mm cell culture dishes and cultivated in 3 mL DMEM at 37 °C and 5% CO₂ until reaching approximately 80% confluency. Cells were then transfected using the polyethylenimine (PEI) method as previously^31^. To ensure robust expression and a balanced ratio between both constructs, 100 ng of pSEMS-aGFPnb-mCherry and 2 µg of pcDNA3.1-mEGFP-ABCA2 DNA were used for transfection. After 6 hours, cells were washed with PBS, detached using trypsin-EDTA, and reseeded onto round glass coverslips (Carl Roth, PK26.2) coated with PLL-PEG-RGD in 3 mL DMEM. Cells were incubated overnight at 37 °C and 5% CO₂. Subsequently, cells were washed three times with prewarmed PBS and fixed with 4% paraformaldehyde (PFA) at 37 °C for 15 minutes. After fixation, cells were washed again with PBS and stored until imaging. On the day of imaging, coverslips were mounted in custom sample holder chambers. Imaging was performed using an inverted confocal laser scanning microscope (IX83-P2ZF, Evident) equipped with two galvanometer-based scanning mirrors. The fluorophores mEGFP and mCherry were excited at 488 nm (OBIS 488 LX, 20 mW; Coherent) and 561 nm (OBIS 561 LS, 20 mW; Coherent), respectively. Fluorescence was collected using a 60× oil immersion objective (PLAPON-SC 60×, NA 1.40; Olympus). Excitation light was reflected by a polychroic mirror (89402bs; Chroma), and emitted fluorescence was filtered using a quadband bandpass emission filter (420–450 nm, 505–530 nm, 578–610 nm, 600–800 nm; 89402m; Chroma). Emission signals were detected sequentially using GaAsP photomultiplier tubes. For acceptor photobleaching experiments, a region of interest (ROI) was defined using the internal stimulation tool of the Olympus acquisition software. mCherry was then bleached using 100% power of the 561 nm laser for 10 seconds.

### Super-resolution DNA PAINT

For DNA-PAINT imaging, HeLa cells were seeded in 35 mm cell culture dishes and cultivated in 3 mL DMEM at 37 °C and 5% CO₂ until reaching approximately 80% confluency. Cells were then transfected using the polyethylenimine (PEI) method with 2 µg of the pcDNA3.1-mEGFP-ABCA2 construct. After 6 hours, cells were washed with PBS, detached using trypsin-EDTA, and reseeded onto round glass coverslips (Carl Roth, PK26.2) coated with PLL-PEG-RGD in 3 mL DMEM, followed by overnight incubation at 37 °C and 5% CO₂ Cells were subsequently washed three times with prewarmed PBS and fixed with 3% paraformaldehyde (PFA) and 0.1% glutaraldehyde at 37 °C for 15 minutes. After fixation, cells were washed with PBS and blocked for 30 minutes in 3% BSA in PBS. Immunostaining was performed overnight at 4 °C using 50 nM anti-GFP nanobody (MASSIVE-TAG-X2 anti-GFP, Massive Photonics GmbH) diluted in antibody staining buffer (PBS supplemented with 3% BSA and 0.1% Triton X-100). The following day, cells were washed three times with PBS and incubated with a 1:5 dilution of 90 nm gold nanoparticles (G-90-20, Absource) as fiducial markers for 5 minutes at room temperature. Samples were then washed three times with PBS and transferred into 1 mL imaging buffer (PBS containing 500 mM NaCl, 1 mM EDTA, and 0.05% Tween-20) supplemented with approximately 100 pM Cy3B-conjugated imager DNA strand. Imaging was performed on a total internal reflection fluorescence microscope (TIRFM) operated in highly inclined and laminated optical sheet (HILO) mode using an inverted microscope frame (IX-81, Olympus), equipped with a motorized quad-line TIR illumination condenser (cellTIRF-4-Line, Olympus) and a motorized xy-stage (Scan IM 120 × 80, Märzhäuser). Three-dimensional single-molecule localization was achieved via astigmatic imaging using a cylindrical lens (Olympus) positioned directly in front of the filter wheel. Transfected cells were identified via GFP fluorescence excited at 488 nm (diode-pumped solid-state laser, max. 150 mW, Olympus), while imager strands were excited at 561 nm (diode-pumped solid-state laser, max. 150 mW, Olympus). Excitation light was delivered through a 100× oil immersion objective (UAPON 100× TIRF, NA 1.49, Olympus). Laser intensities for imager strand excitation were typically adjusted to ∼30 W/cm². Fluorescence emission was filtered using bandpass filters (BrightLine HC 525/50 for GFP and BrightLine HC 600/37 for Cy3B, Semrock) and detected using an sCMOS camera (ORCA-Flash4.0 V3, Hamamatsu). Image acquisition was controlled using cellSens 2.2 software (Olympus), recording 80,000 frames with an exposure time of 50 ms and 2×2 pixel binning, resulting in an effective pixel size of 130 nm. During acquisition, the focal plane was stabilized using a hardware autofocus system (IX2-ZDC2, Olympus). The temperature was maintained at 25 °C using a large incubation chamber (TempController 2000–2, Pecon), while sample humidity was controlled using a stage-top incubator (CO₂-Controller 2000, Pecon) to prevent buffer evaporation. Axial calibration for astigmatic point spread functions (PSFs), required for 3D localization, was performed at the beginning of each imaging session by acquiring z-stacks of immobilized fluorescent TetraSpeck™ microspheres (100 nm diameter, Invitrogen, T7279). Stacks were recorded in imaging buffer with a step size of 10 nm using a piezo z-stage (NanoScanZ, NZ100, Prior Scientific).

### DNA-PAINT post-processing

Raw datasets were processed using the Picasso software package^32^ (https://github.com/jungmannlab/picasso). First, the calibration z-stack of TetraSpeck™ beads was analyzed using Picasso Localize. For single-bead identification, the box size was set to 13 pixels, and the minimum net gradient threshold was adjusted to suppress weak background localizations. Photon conversion parameters were set as follows: EM gain = 1, baseline = 400, sensitivity = 0.46, quantum efficiency = 0.80, and pixel size = 130 nm. A calibration file for 3D localization was generated using the Calibrate 3D function in Picasso Localize. The same photon conversion parameters and the generated calibration file were subsequently applied to the raw sample datasets. The minimum net gradient threshold was further adjusted to exclude nonspecifically bound imager strands and background signals, thereby enriching for high-confidence localizations. Single-molecule events were fitted using a least-squares Gaussian model. For 3D localization, a magnification factor of 0.7 was applied. Processed datasets were then imported into Picasso Render for visualization and post-processing. Drift correction was first performed using cross-correlation, followed by a second correction step based on fiducial markers (gold nanoparticles).

## Results

### Structural and functional characterization of full-length human ABCA2

To investigate the functional role of ABCA2 in lysosomal lipid homeostasis, we performed single-particle cryo-EM analysis of ABCA2 in the presence and absence of cholesteryl hemisuccinate (CHS) (Extended Data Fig. 1 and 2). We overexpressed full-length human ABCA2 in HEK293S GnTI^-^ cells and purified it in either LMNG or LMNG/CHS (Extended Data Fig. 3a). The ATPase activity of both preparations was quantified using an enzyme-coupled ATPase assay, yielding hydrolysis rates approximately sevenfold higher in the CHS-supplemented preparation (112 nmol/min/mg) than in the CHS-free preparation (17 nmol/min/mg) (Extended Data Fig. 3b). Our five high-resolution ABCA2 structures reveal the conformational transitions associated with cholesterol transport from the TMDs to the ECDs (Fig. 1a). Variations in their respective inter-NBD distances enabled a clear assignment of the transport cycle sequence, which we identify as the apo (nucleotide-free state), closed and return conformations (ADP-bound post-hydrolysis states) (Fig. 1b).

**Fig. 1.**
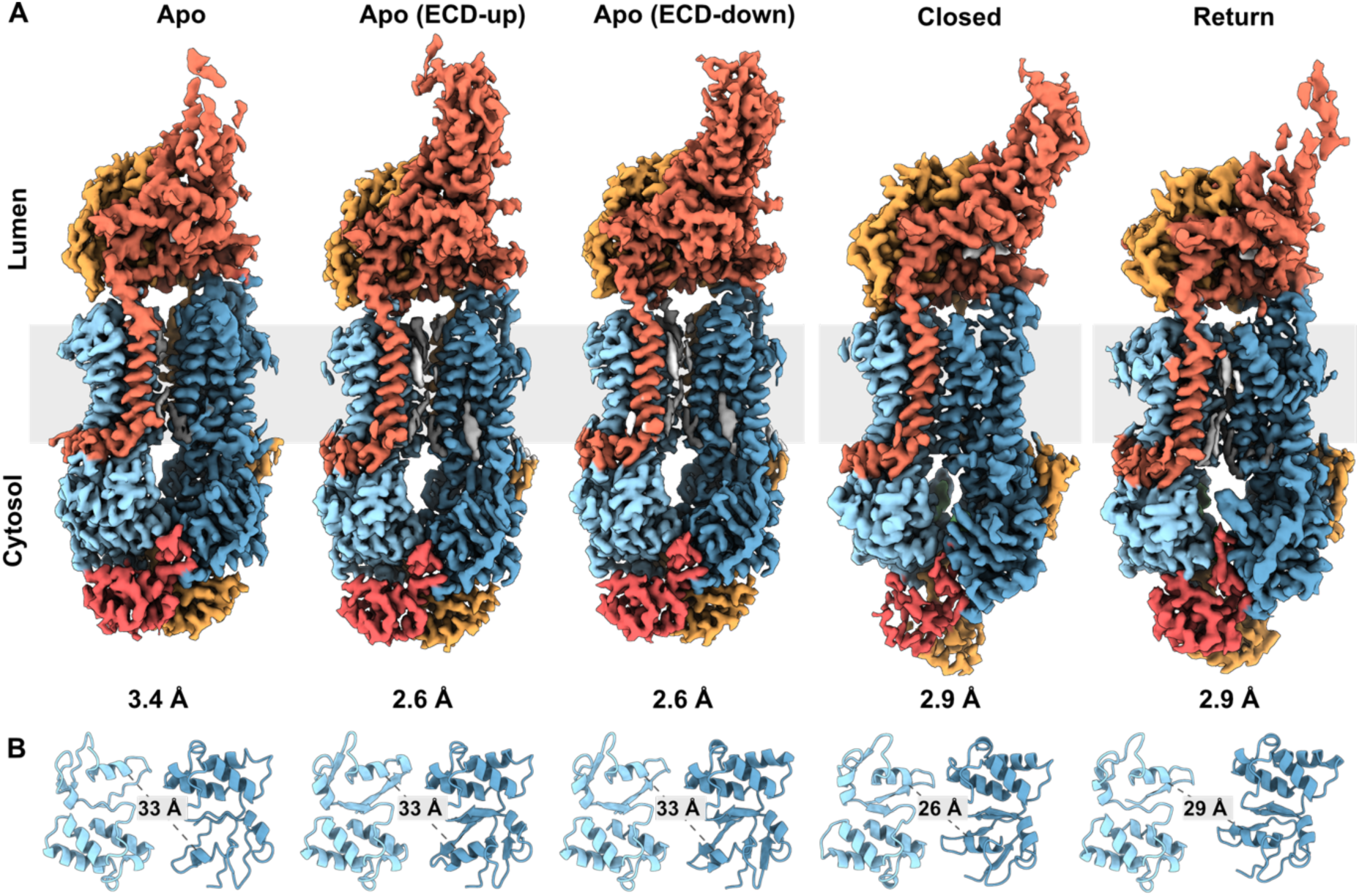
ABCA2 in five transport intermediates. **(A)** EM densities are displayed in membrane orientation and colored to highlight the domain architecture (salmon: TM1 and ECD1; light blue: TM2-TM8 and NBD1; mustard: RD1, TM9 and ECD2; dark blue: TM10-TM16; red: RD2). Gray boxes indicate the approximate membrane boundaries. **(B)** NBDs of ABCA2 corresponding to conformations in (A) viewed from the cytoplasm, shown in cartoon representation. Distance measurements between the NBDs of all transport states define their conformational sequence.

We confirmed the presence of ABCA2 in lysosomes through re-analysis of a lysosomal pull-down mass spectrometry experiment^33^ (Extended Data Fig. 3c). Additionally, to define the orientation of ABCA2 within the intracellular organelle, we co-expressed mEGFP-tagged ABCA2 and a cytosolically expressed anti-GFP nanobody fused to mCherry in HeLa cells, followed by analysis using dual-color confocal fluorescence microscopy. Binding of the cytosolically expressed anti-GFP nanobody fused to mCherry clarified that the NBDs of ABCA2 are oriented towards the cytosol (Extended Data Fig. 3d, e).

### ABCA2 in the nucleotide-free state

Our three high-resolution nucleotide-free structures, one purified in LMNG at 3.4 Å and two in LMNG/CHS at 2.6 Å each, jointly provide novel insights into the overall architecture of the transporter (Fig. 2a). Delipidation and relipidation of ABCA2 purified from LMNG and LMNG-CHS enabled us to distinguish endogenous lipid-binding sites, while the overall structure remained similar despite changes in the purification environment. Compared to published structures of ABCA transporters, ABCA2 exhibits pronounced differences in the arrangement of individual domains; particularly, the observed concerted rotations of the ECDs and RDs have not yet been described for other ABCA transporters. Both ECDs are interconnected through two shared six-bladed beta sheets and stabilized by four disulfide bonds: one interdomain bond between C694 and C1643, one within ECD1 (C75-C418), and two within ECD2 (C1543-C1576, C1652-C1641) (Fig. 2c). The ECDs extend approximately 90 Å into the lysosomal lumen and are heavily glycosylated. Our maps showed densities for 11 of the 25 distinct glycosylation sites predicted by Uniprot, which are important for the stability and folding of ABC-A family members^34^ (Supplementary Fig. 1). On the opposite side of the membrane leaflet, the cytoplasmic NBDs and RDs extend approximately 60 Å from the membrane. The TMDs of ABCA2 consist of two pseudo-symmetrical halves, each composed of six full transmembrane helices (TM1-5 and 8, TM9-13 and 16) and one reentrant helix (TM6-7 and TM14-15). Like the ABCG subfamily, ABCA transporters feature a non-swapped architecture in which each TMD remains within its respective half, and the NBDs are positioned close to each other in the nucleotide-free state.

Tracing the polypeptide chain from the N-terminus, a cytoplasmic elbow helix leads into TM1, which traverses the lipid bilayer, crosses the pseudo-symmetrical axis, and enters the larger luminal ECD1. Within ECD1, TM1 contributes directly to the formation of the first six-bladed beta-sheet of mixed orientation, which connects ECD1 and ECD2. The second blade is oriented antiparallel to the first and extends into TM2 within the lipid bilayer. A four-helix bundle of antiparallel alpha helices is positioned between the first and second antiparallel blades of ECD1, extending perpendicular to the membrane and forming a hollow hydrophobic vertical tunnel that is filled with cholesterol molecules and lipid. A further globular domain of ECD1, which extends from the loops of the helix bundle, could not be resolved in our EM density. ECD1 also contributes four blades to the second shared beta-sheet, which exhibits the same topology as the first. In analogy, TM9, the first transmembrane helix of the second half-TMD, leads into ECD2 and immediately contributes to the first blade of the mixed beta-sheet that ties the ECDs together, as described above. The final blade of ECD2 forms part of a second mixed-orientation bladed beta-sheet that is located opposite the first six-bladed beta-sheet. As such, the final blade runs parallel to the beta-sheet from ECD1 and leads into TM10. Structural analysis of ABCA2 under reducing conditions resulted in highly destabilized ECDs, while the overall architecture remained intact (Extended Data Fig. 4a). Curiously, destabilization of the ECDs had a pronounced effect on ATPase activity, as indicated by reduced ATP hydrolysis (Extended Data Fig. 4b), highlighting direct crosstalk between the ECDs and NBDs/RDs, and underscoring the importance of intact ECDs for full functionality.

**Fig 2.**
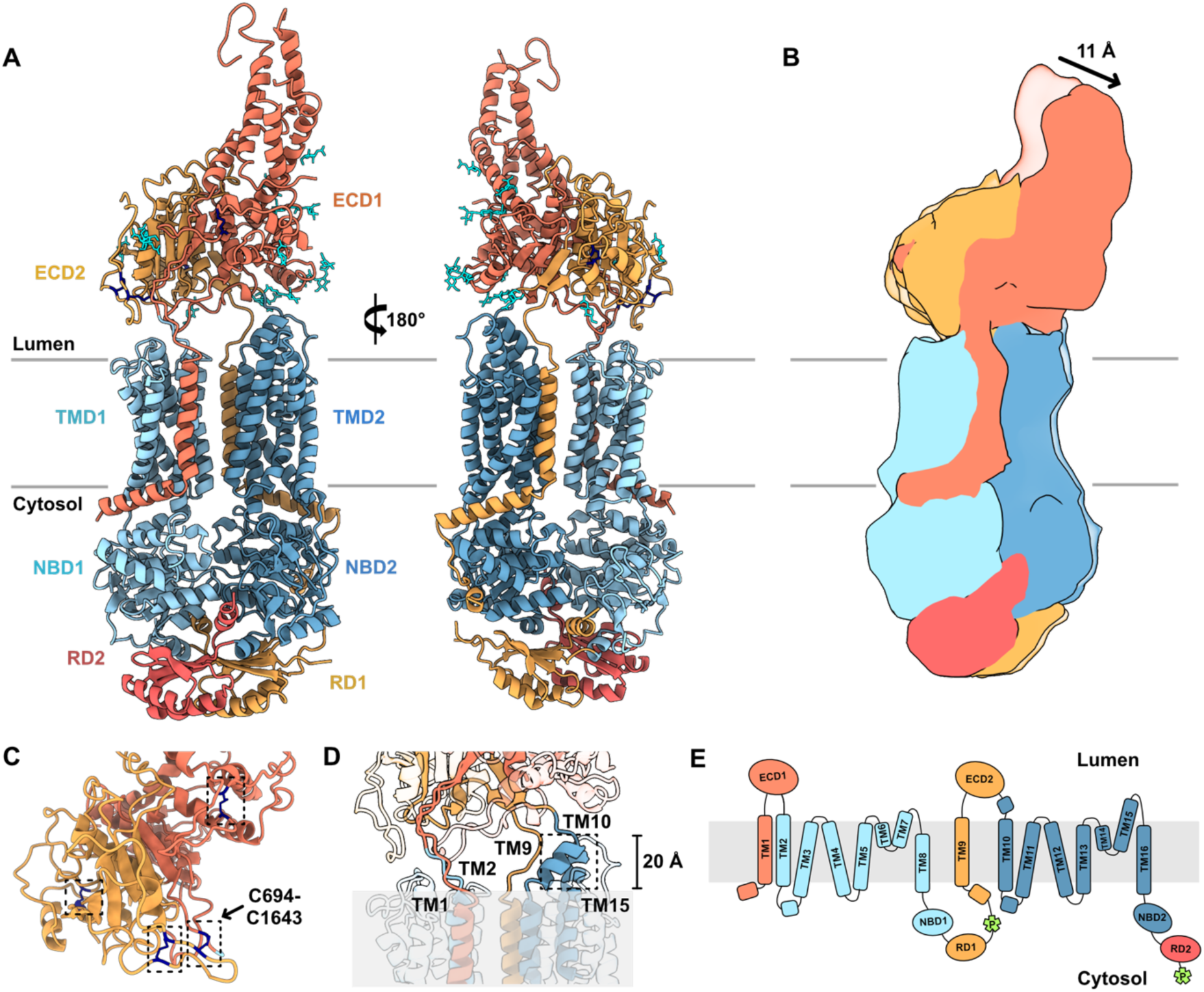
Overall architecture of human ABCA2 in the nucleotide-free state. Cryo-EM structure of ABCA2 in two orthogonal views in cartoon representation. **(A)** The structure reveals the canonical domain organization of ABCA-family transporters, consisting of a pair of TMDs, NBDs, ECDs and RDs arranged in a pseudo-symmetrical architecture. Individual structural domains are color-coded as in Figure 1; the first helix of TMD1 and ECD1 in salmon, ECD2 and RD1 in yellow, TMD1 and NBD1 in light blue, TMD2 and NBD2 in dark blue, and RD2 in red. The disulfide bonds in the ECDs are shown in dark blue, and the N-acetylglucosamine glycosylations in cyan. **(B)** Comparison of conformational differences within the ECDs between two nucleotide-free states (light grey: apo ECD-up; colored: apo ECD-down). **(C)** Disulfide bond (dark blue) distribution in the ECDs, with C694-C1643 corresponding to an interdomain bond. **(D)** TM1, TM2, TM9 and TM10 connect ECDs to the remaining part of the protein. TM15 and the kinked helix TM10 extend approximately 20 Å from the membrane. **(E)** Topological cartoon of ABCA2. Lime green square star shapes represent phosphorylation sites.

Within TMD1, the coupling helix between TM2 and TM3 engages with NBD1. As in other A-family ABC transporters, TM6 and TM7 form reentrant helices that extend halfway into the membrane bilayer. A long loop connects the C-terminus of TM7 to TM8, and from TM8, another extended, partially resolved loop connects to NBD1. In TMD2, the coupling helix between TM10 and TM11 engages NBD2. The architecture of TMD2 resembles TMD1, except that the reentrant helices in TMD2 (TM14 and TM15) are elongated and display additional helical turns. Likewise, TM10, which connects to ECD2 and the coupling helix, is lengthened by a kinked helix protruding from the membrane and is absent in TMD1. Accordingly, helices TM10 and TM15 each extend approximately 20 Å from the luminal leaflet of the membrane (Fig. 2d). In the apo states, the TMDs are oriented toward the luminal side of the lipid bilayer and exhibit a 20 Å-wide lateral cleft spanning the bilayer, a structural feature not accurately predicted by AlphaFold (Extended Data Fig. 4c).

The NBDs follow the classical fold and the high structural conservation within the ABC transporter family, including canonical ATP-binding sites. The RDs emerge from the zipper helices, engage in domain swapping, and are anchored to the base of their respective opposite NBDs. Furthermore, the RDs host two phosphorylation sites (Fig. 2e), which may be associated with NBDs dimerization^35^. Each RD consists of two adjacent and intersecting beta hairpins, which are succeeded by a short, negatively charged alpha helix, sandwiched between both NBDs at the positively charged region. Following RD1, a long loop extends along the vertical side of NBD2, where the first phosphorylation site (S1327-S1331) is located, and connects to the elbow helix TM9 of the second pseudo-symmetric half of the transporter. The second half follows a similar structural path and leads to RD2 at the C-terminus, where the second phosphorylation site (S2412-T2419) is situated in a long loop that was not resolved in our density.

Notably, we identified two distinct conformations in the nucleotide-free state (apo ECD-up and apo ECD-down), with conformational differences primarily in the ECDs. The ECDs swivel approximately 11 Å between the two conformations, indicating conformational flexibility at the ECDs-TMDs interface, while the overall structure of the TMDs, NBDs, and RDs remains unchanged (Fig. 2b). The swiveling of the ECDs coincides with movement of TM1, TM2, TM9, and TM10, which connect the ECDs to the rest of the protein.

### Conformational changes of ABCA2 after ATP hydrolysis

**Fig 3.**
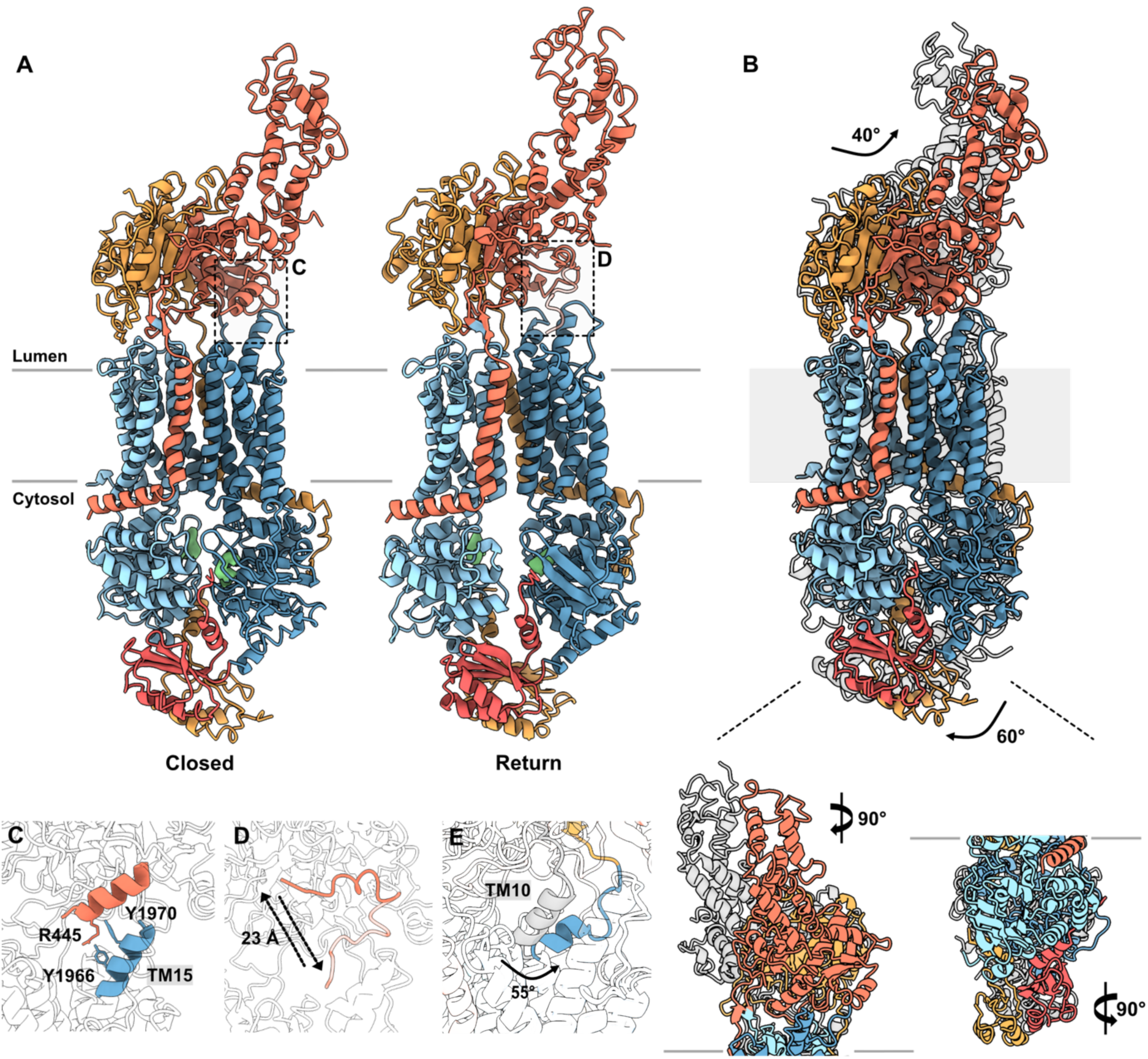
Closed and return conformations of ABCA2. **(A)** Cryo-EM structures of ABCA2 in ADP-bound post-hydrolysis states, closed and return conformations. Domains are colored as in Fig. 1. The black dotted box in the closed conformation highlights the interaction between helix residues 440-455 and reentrant helix TM15. In the return conformation, the black dotted box highlights the movement of the same helix residues 440-455 within the same structure. **(B)** Comparison of conformational differences between the apo and closed conformations in different views, showing a 40° anticlockwise rotation of the ECDs and a 60° clockwise rotation of the RDs. **(C)** The interaction between the helix and TM15 at R445 with Y1966 and Y1970, depicted using the closed state as an example. **(D)** Close-up view of the helix movement within ECD1 between the return conformations, exhibiting a vertical movement of 23 Å. **(E)** Close-up view of the apo and closed conformations showing a 55° swiveling movement of TM10.

To investigate the transport mechanism of ABCA2, we conducted cryo-EM analysis in the nucleotide-bound state after incubation with MgATP and orthovanadate. Although we intended to capture a hydrolysis transition state, the resulting maps revealed clear ADP densities at both nucleotide-binding sites, which are not fully dimerized (Fig 3a, d). A similar scenario is observed for ABCA7, where the addition of ATP*γ*S did not induce NBD dimerization^36^, suggesting a common effect. In the ADP-bound post-hydrolysis state, we resolved two distinct conformations of ABCA2, each at 2.9 Å resolution. In one, the TMDs are fully closed with more tightly associated NBDs, comparable to the nucleotide-bound structures of other ABCA transporters ^36–40^. We refer to this as the closed conformation. A second conformation features partially separated TMDs and NBDs, which we refer to as the return conformation. In this density map, we identified two alternative conformations of a short helix (residues 440-455) within ECD1 at the TMD interface, with the helix shifting by approximately 23 Å (Fig. 3d). In the closed conformation, the helix is located between the two alternative helix positions observed in the return conformation, where R445 interacts with Y1966 and Y1970 of TM15 (Fig. 3c).

In both closed and return structures, major interconnected conformational changes are observed across all domains. In the closed conformation, the TMDs pivoted around the N-terminal regions of TM5 and TM11 as a rigid body, causing complete collapse of the lateral pocket. This is accompanied by a previously unreported reorientation of the RDs, which rotate horizontally by approximately 60° relative to their position in the apo conformation (Fig. 3b). The extended kinked helix of TM10 swivels backward by approximately 55° (Fig. 3e) and, together with TM9, mediates RDs repositioning toward the ECDs, resulting in a dramatic rotation of the ECDs by approximately 40° orthogonal to the membrane interface (Fig. 3b). In the return conformation, the lateral pocket is slightly open and adopts an inverted V-shape. Additionally, the NBDs gradually separate, with 30 Å compared to 26 Å in the closed conformation, as shown in Figure 1c. Despite these significant rearrangements of the TMDs and NBDs, both ECDs and RDs maintain their positions between the closed and return conformations (Fig. 3a).

### Lipid transport sites of ABCA2

**Fig 4.**
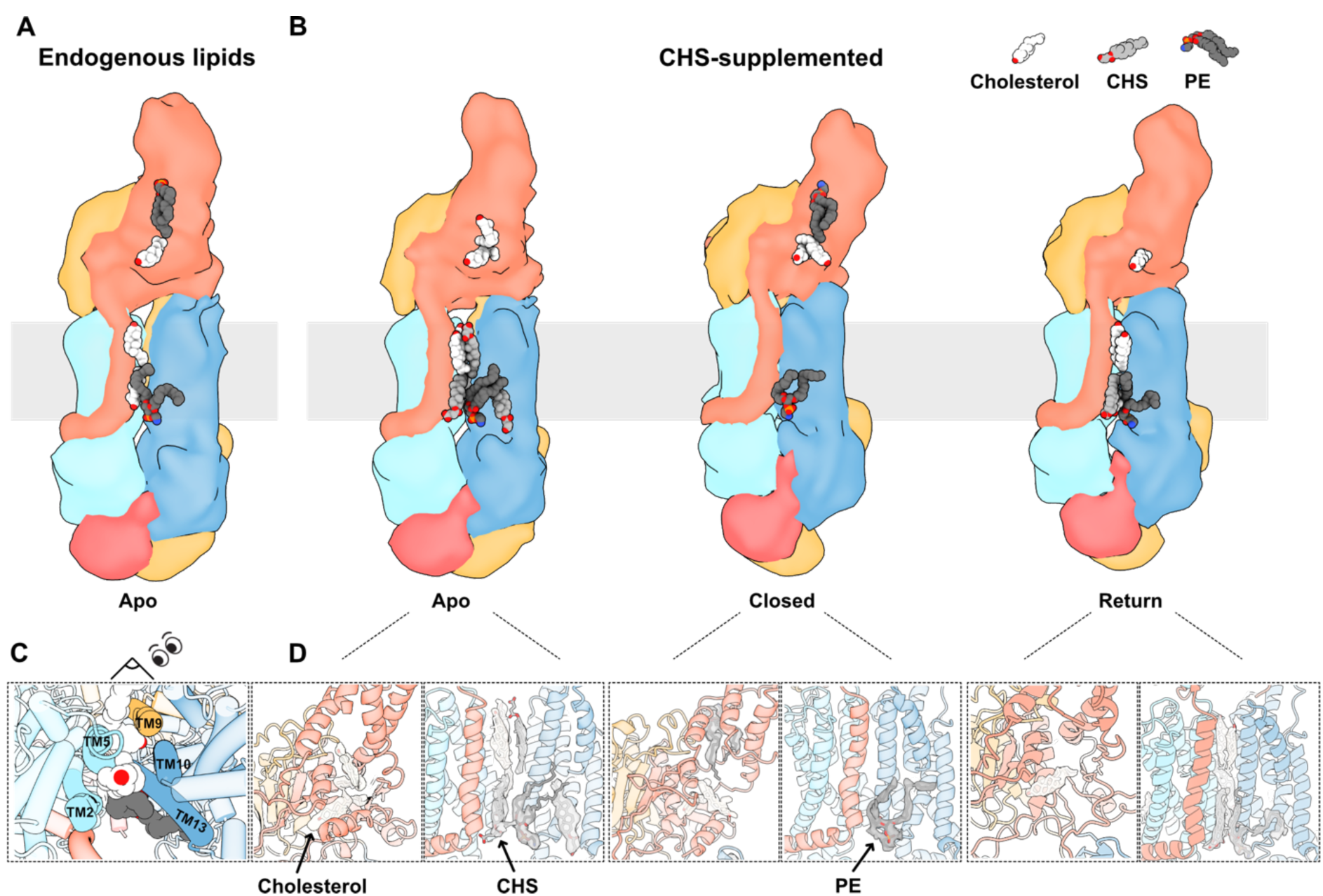
Cholesterol, CHS, and phospholipid binding sites in ABCA2 across all conformational states. **(A)** Apo conformation of ABCA2 from a CHS-free preparation in cartoon representation colored as in Fig. 1, with endogenous lipids shown as spheres with cholesterol in white and PE in dark grey. **(B)** Surface representation of the apo, closed, and return conformations from a CHS-supplemented preparation, with cholesterol and PE as the same color as in (A) and CHS in light grey, shown in sphere representation. **(C)** TMDs of ABCA2 corresponding to conformation in (A) viewed from the lumen, showing the cholesterol transport site. **(D)** Close-up view of cholesterol, CHS and PE densities within the TMDs and ECDs of all conformational states.

All our structures exhibit multiple lipid densities, mostly corresponding to either endogenous cholesterol or CHS, which we could distinguish by comparing samples prepared with and without CHS supplementation (Fig. 4a, b and Extended Data Fig. 5a). Most cholesterol densities are located between TM2, TM5, TM9, TM10 and TM13, forming a lateral lipid-binding pocket spanning both membrane leaflets (Fig. 4c). Additionally, we observed either one or two cholesterol densities within ECD1 across all structures (Fig. 4a, d). These cholesterol molecules are closely positioned within a hydrophobic tunnel, forming a queue from the bottom of ECD1. In all structures, a cholesterol molecule resides within ECD1 near helix 440-455. Additionally, we detected an extra density near the lid of ECD1 and adjacent to the lateral binding pocket of the TMD, resembling a phosphatidylethanolamine (PE) lipid. Notwithstanding, we also identified several CHS densities near the lipid-binding pocket of the TMD and surrounding the protein within the membrane bilayer in the CHS-supplemented preparation.

## Discussion

**Fig 5.**
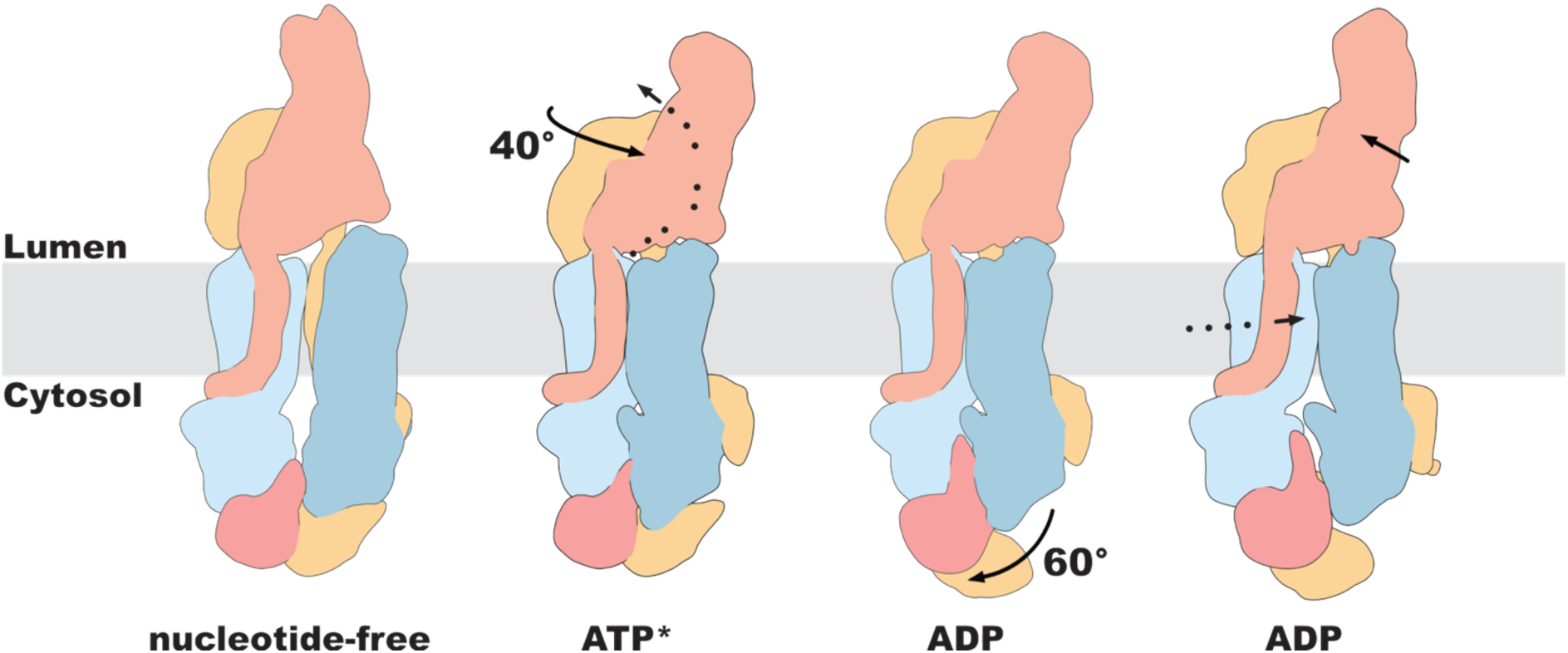
Representative mechanism of ABCA2 in nucleotide-free and ADP-bound post-hydrolysis states delineating the cholesterol transport pathway on the lysosomal membrane. Dotted arrows indicate cholesterol translocation; solid lines indicate individual domain rotations.

The distinct conformations delineate the cholesterol transport mechanism of ABCA2 and highlight unique features of the transporter (Fig. 5). In the nucleotide-free state, the transporter opens longitudinally only to one side, facilitating lipid and cholesterol access from both membrane leaflets. In the closed conformation, ATP binding induces NBD dimerization and a pronounced rotation of the ECD and RD. The tightly dimerized TMDs no longer permit lipid retention, indicating lipid sequestration towards the ECDs. Reminiscent of transporters of the B-family^41^, in the subsequent return conformation, the TMDs and NBDs open progressively into an inverted V-shaped TMD arrangement that allows lipid loading from the membrane bilayer.

Our structures reveal several features not observed in other ABCA family members. Most ABCA transporters, including ABCA1, ABCA3, and ABCA4 in various hydrophobic environments, as well as ABCA7 in detergent, exhibit a characteristic V-shaped lateral opening of the TMDs in the pre-state^37,38,42^. In these structures, the N-termini of TM5 and TM11, which face the lower membrane leaflet, are positioned in proximity, allowing direct side chain interactions, while the C-termini are widely separated. This fenestration between the TMDs may facilitate horizontal lipid access from either side of the transporter. The only exception to this fenestration is the structure of ABCA7 in a nanodisc (PDB IDs: 8DEW, 8EE6), which displays a wide, parallel separation between the respective TMD halves. Our apo conformation closely resembles the fenestration of ABCA7, where TM5 and TM11 are oriented parallel to each other and orthogonal to the membrane. However, in ABCA2, the fenestration is closed on one side, resulting in a narrow opening on the opposite side and forming a distinct lipid-filled pocket accessible only from one side of the transporter.

A structural comparison of the two distinct conformations in the post-hydrolysis state reveals a previously uncharacterized intermediate: the return conformation. In this conformation, the unique inverted V-shape at the lateral lipid-binding pocket may facilitate lipid loading from the membrane to the TMDs before adopting a parallel opening in the apo state. The conformational transition from a collapsed lateral pocket to an inverted V-shaped TMD cavity may induce membrane curvature, mediating interactions between residues within the hydrophobic cleft of the TMDs and the surrounding lipids in the bilayer^43^, thereby enabling lipid substrate loading to the transporter.

Our structures reveal large-scale interdependent rotations of both ECDs and RDs in the closed and return conformations, a phenomenon not observed in other transporters and suggest a mechanism for lipid loading and regulation of the transport cycle. The rotational movement of the ECDs and RDs is most apparent in the closed state. Here, the TMDs resemble a configuration previously reported for other ABCA family members; however, in our structure, the RDs exhibit a pronounced anticlockwise rotation of 60°. We suggest that, in the cellular environment, phosphorylation of the RDs may function as an unlocking mechanism that permits NBD dimerization upon ATP binding^44^, which may explain an increase in negative charges in the RD region upon rotation (Extended Data Fig. 5b). The extended kinked helix of TM10 (Fig. 3e), a feature uniquely shared by ABCA2 and ABCA3, is likely to play a primary role in the swiveling movement of the ECDs. In contrast, TM1, TM2, and TM9, which are also connected to the ECDs, do not rotate in concert with the RD. Although a similar extended kinked helix is observed in ABCA3, ABCA3 does not exhibit a pronounced RD rotation compared to ABCA2^38^. Correspondingly, the ECD rotation in ABCA2 between the nucleotide-free and nucleotide-bound states resembles the movement observed in ABCA4, but ABCA4 does not exhibit RD rotation^42^.

In the CHS-supplemented preparation of the nucleotide-free state, lipids assemble in two concentric layers within the lipid-binding pocket of the TMDs. The outer layer contains CHS molecules and a phospholipid identified as phosphatidylethanolamine (PE). Peculiarly, the inner binding pocket contains only endogenous cholesterol molecules stacked in a single column between the upper and lower leaflets, similar to the CHS-free preparation. Such positioning and vertical orientation have not been observed in other ABCA transporters but suggest a mechanism for lipid extraction prior to entry into the ECDs. In analogy to studies on ABC transporters of the B-family, we attribute the higher ATPase activity of the CHS-containing preparation compared with the CHS-free preparation to the purification environment, which strongly influences transporter activity^45,46^. Additionally, the supplementation with CHS may stimulate ABCA2 activity as a transport substrate, consistent with its function as a lipid transporter.

Dimerization of the TMDs results in the sequestration of all bound lipids, including cholesterol and likely PE, into a molecular basket, before entering the ECDs. Consistent with previously resolved ABCA transporter structures showing phospholipids^42,47^, we identified a PE lipid at the TM5/11 interface, which resides in a similar position throughout the transport cycle. Another PE lipid was also observed near the lid of ECD1 in all conformations except the return conformation, suggesting that ABCA2 could transport other lipids besides cholesterol. In fully loaded ECDs, we identified three lipid-binding positions, while all conformations exhibited a single conserved cholesterol-binding site not previously reported in any ABCA transporter (Fig. 4a, b and d). Interestingly, this cholesterol-binding site is located adjacent to helix 440-455 within ECD1, which appears to be important for anchoring to reentrant helix 15 via cation-π interactions, thereby drawing the ECDs toward the TMDs. This interaction may facilitate the movement of cholesterol molecules into the hydrophobic tunnel of the ECD1 (Extended Data Fig. 5c). Cholesterol may also contribute to the stabilization of the ECDs and to the interaction between helix 440-455 and reentrant helix 15.

Unlike other ABCA transporters, ABCA2 is predominantly expressed in the brain, particularly in oligodendrocytes, which are responsible for synthesizing most of the cholesterol required for myelination^1,48^. Therefore, proper folding of the ECDs is crucial for ABCA2 function, and mutations in cysteine residues within the ECDs may lead to dysregulated lipid metabolism in oligodendrocytes. Within the ECDs, cholesterol traverses the hydrophobic tunnel and exits via the ECD1 lid (Extended Data Fig. 6). Although the ECDs of ABCA1 have been shown to bind the extracellular protein apolipoprotein A, there is currently no evidence that the ECDs of ABCA2 bind to a lipid acceptor. Alternatively, ABCA2 may facilitate the formation of lipid-rich lamellae within lysosomes, as characteristic in oligodendrocytes^49^. These lipid-rich lamellar lysosomes could subsequently fuse with the plasma membrane to release lipids, thereby supporting myelin sheath formation. A similar phenomenon is observed in both ABCA3 and ABCA12, in which lamellar bodies fuse with the plasma membrane to secrete lipids into the alveolar lumen and the skin barrier, respectively^34,50^. Notably, ABCA2 has also been reported to localize to the limiting membrane of lamellar bodies in type II alveolar cells^51^, where it may fulfill a similar function. Our study establishes a molecular basis for understanding how ABCA2 mediates cholesterol transport across the lysosomal membrane in oligodendrocytes and provides further insight into ABCA2’s role in myelin sheath biogenesis.

## Author contributions

S.M.T. and A.M. designed and conducted the study with help from K.P. and D.J.. S.M.T performed the biochemical and cryo-EM sample preparation with help from N.V., M.N., D.S., K.P. and D.J.. S.M.T. and K.S. performed the single-particle analysis. S.M.T. and K.P. built the atomic models. B.E. and F.F. contributed mass spectrometry data. M.H. and J.P. contributed all light microscopy. S.M.T. and A.M. wrote the original draft. All authors discussed the data and edited the final version. A.M. secured funding for this study.

## Acknowledgements

We thank Hikade, H. and Huber L. for technical support. This work was supported by a grant of the DFG (MO2752/4-1), the SFB 1557, the DFG INST190/196-1 FUGG and the RTG 2900 (all A.M.).

## Data availability

All density maps and models have been deposited in the Electron Microscopy Data Bank and the PDB.

